# Robust and conserved stochastic self-assembly mechanism for dynamic ParB-*parS* partition complexes on bacterial chromosomes and plasmids

**DOI:** 10.1101/345066

**Authors:** Roxanne Diaz, Aurore Sanchez, Jérôme Rech, Delphine Labourdette, Jérôme Dorignac, Frédéric Geniet, John Palmeri, Andrea Parmeggiani, François Boudsocq, Véronique Anton Leberre, Jean-Charles Walter, Jean-Yves Bouet

## Abstract

Chromosome and plasmid segregation in bacteria are mostly driven by ParABS systems. These DNA partitioning machineries rely on large nucleoprotein complexes assembled on centromere sites (*parS*). However, the mechanism of how a few *parS*-bound ParB proteins nucleate the formation of highly concentrated ParB clusters remains unclear despite several proposed physico-mathematical models. We discriminated between these different models by varying some key parameters *in vivo* using the plasmid F partition system. We found that ‘Nucleation & caging’ is the only coherent model recapitulating *in vivo* data. We also showed that the stochastic self-assembly of partition complexes (i) does not directly involve ParA, (ii) results in a dynamic structure of discrete size independent of ParB concentration, and (iii) is not perturbed by active transcription but is by protein complexes. We refined the ‘Nucleation & Caging’ model and successfully applied it to the chromosomally-encoded Par system of *Vibrio cholerae*, indicating that this stochastic self-assembly mechanism is widely conserved from plasmids to chromosomes.

## Introduction

The segregation of DNA is an essential process for the faithful inheritance of genetic material. Minimalistic active partition systems, termed Par, ensure this key cell cycle step in bacteria (Baxter and Funnell, 2014) and archaea (Schumacher et al., 2015). Three main types of bacterial partition systems have been identified and classified by their NTPase signatures. Of these, the type I, also called ParAB*S*, is the only one present on chromosomes and the most widespread on low-copy number plasmids (Gerdes et al., 2000). Each replicon encodes its own ParAB*S* system and their proper intracellular positioning depends on the interactions of the three ParAB*S* components: ParA, a Walker A cytoskeletal ATPase; ParB, a dimer DNA binding protein; and *parS*, a centromere-like DNA sequence that ParB binds specifically. The ParA-driven mechanism that ensures the proper location and the directed segregation of replicons relies on the positioning of ParB*S* partition complexes within the nucleoid volume (Le Gall et al., 2016) and on a reaction diffusion-based mechanism (Hu et al., 2017; Hwang et al., 2013; Lim et al., 2014; Walter et al., 2017).

The centromere-like *parS* sites are located close to the replication origin on chromosomes and plasmids, and are typically composed of 16-bp palindromic motifs (Lin and Grossman, 1998; Mori et al., 1986). ParB binds with high affinity to its cognate *parS* as dimers (Bouet et al., 2000; Hanai et al., 1996). This serves as a nucleation point for assembling high molecular weight ParB -*parS* partition complexes, as initially seen by the silencing of genes present in the vicinity of *parS* (Lobocka and Yarmolinsky, 1996; Lynch and Wang, 1995). ParB binds over 10-Kbp away from *parS* sites for all ParAB*S* systems studied to date (Donczew et al., 2016; Lagage et al., 2016; Murray et al., 2006; Rodionov et al., 1999; Sanchez et al., 2015). This phenomenon, termed spreading, refers to the binding of ParB to centromere-flanking DNA regions in a non-specific manner. The propagation of ParB on DNA adjacent to *parS* is blocked by nucleoprotein complexes such as replication initiator complexes in the case of the P1 and F plasmids (Rodionov et al., 1999; Sanchez et al., 2015), or repressor-operator complexes on the bacterial chromosome (Murray et al., 2006). These ‘roadblock’ effects led to the initial proposal that ParB propagates uni-dimensionally on both sides of the *parS* sites, in a so-called ‘1D-spreading’ model. However, this model was put into question as (i) the quantity of ParB dimers present in the cell was insufficient to continuously cover the observed spreading zone, and (ii) ParB binding to *parS* adjacent DNA resisted biochemical demonstration (reviewed in Funnell, 2016).

As an alternative to ‘1D-spreading’, two other models for partition complex assembly have been proposed, namely ‘Spreading & bridging’ (Broedersz et al., 2014) and ‘Nucleation & caging’ (Sanchez et al., 2015). Both models rely on strong ParB clustering with over 90% of ParB confined around *parS* (Sanchez et al., 2015). The ‘Spreading & bridging’ model proposes that nearest neighbour interactions (1D-spreading) initiated at *parS* and non-*parS* DNA sites in combination with their subsequent interactions in space (3D-bridging), lead in one of the conditions tested (strong spreading and bridging) to the condensation of the ParB-bound DNA into a large 3D complex over a contiguous 1D DNA domain (Broedersz et al., 2014; Graham et al., 2014). The ‘Nucleation & caging’ model rather proposes that the combination of dynamic but synergistic interactions, ParB-ParB and ParB-nsDNA (Fisher et al., 2017; Sanchez et al., 2015), clusters most of the ParB around *parS* nucleation sites where a few ParB dimers are stably bound (Fig. 1A). The *in vivo* ParB binding pattern from high resolution ChIP-sequencing data was described with an asymptotic decay as a characteristic power-law with an exponent b= −3/2, corresponding to the decreasing probability of the DNA to interact with the ParB cluster as a function of the genomic distance from *parS* (Sanchez et al., 2015). This model therefore proposes that the DNA surrounding the *parS* site interacts stochastically with the sphere of high ParB concentration. Interestingly, these three different assembly mechanisms have been explicitly modelled (Broedersz et al., 2014; Sanchez et al., 2015), thus allowing their predictions to be experimentally tested.

**Figure 1.**
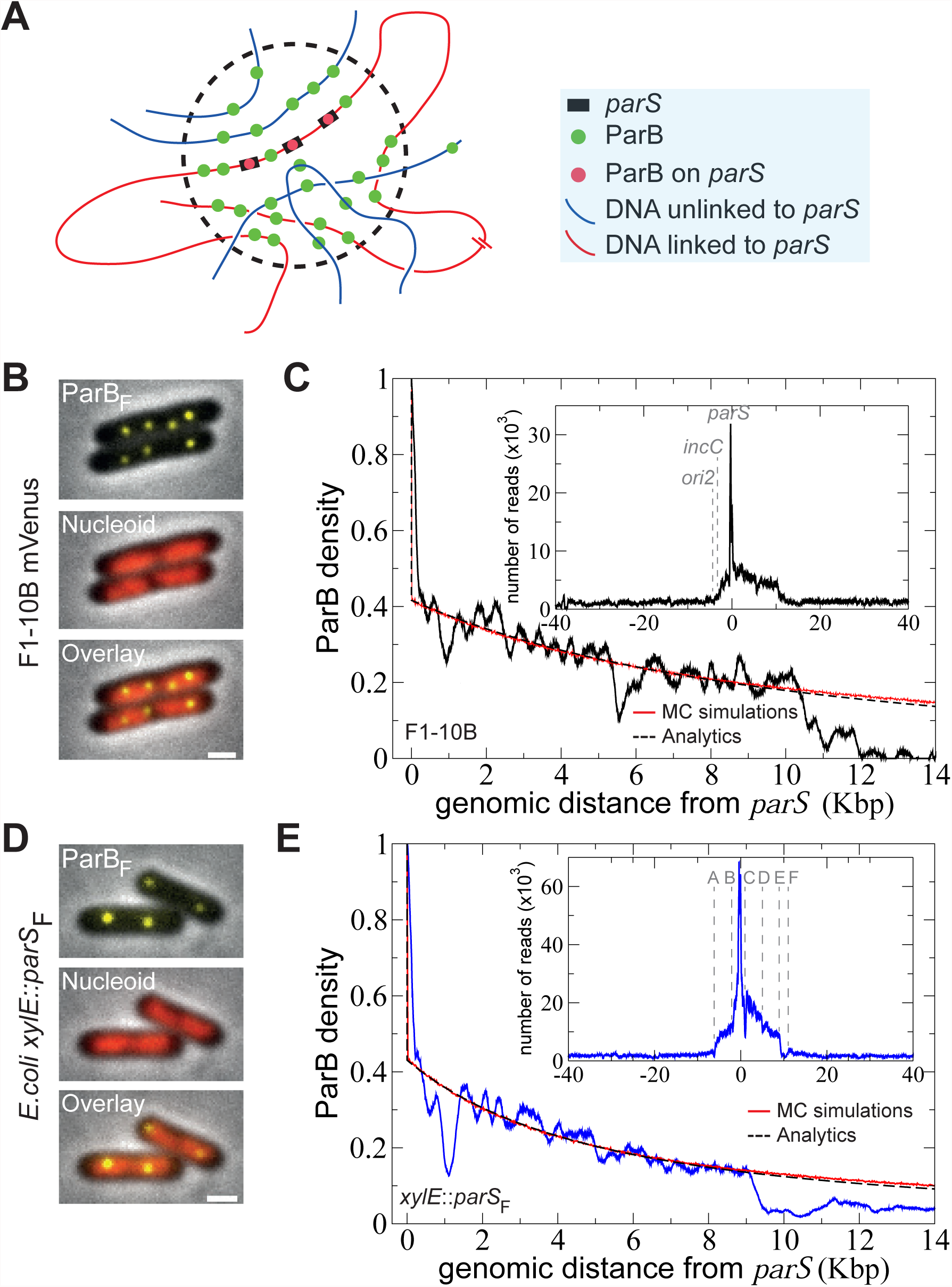
ParB_F_ binding outside of *parS* centromere on plasmid and chromosome. (A) Schematic representation of the ‘Nucleation & caging’ model. Most ParB dimers (green dots) are highly confined in a cluster (dotted circle) centered on the *parS* sites (black rectangles) onto which some ParB are stably bound (red dots). The DNA entering the cluster is bound stochastically by ParB. Red and blue lines represent DNA present at small and large (or on a different molecule) genomic distance from *parS*, respectively. (B) ParB clusters on plasmid F *in vivo*. Typical *E. coli* cells (DLT3594) displays foci of ParB_F_-mVenus protein (top) expressed from the endogenous genetic locus of the plasmid F (F1-10B-mVenus). The nucleoid is labelled with Hu-mCherry (central). The overlay (bottom) combines the two fluorescent channels. Over 99 % of cells harbor ParB_F_ foci. White bars: 1 µm. (C) ParB_F_ binding outside *parS*_F_ on the plasmid F follows a power-law decay. High resolution ChIP-seq performed on DLT3586 carrying the plasmid F (F1-10B). The ParB density, normalized to 1 at the first bp after the last *parS*_F_ binding repeat, is displayed over 14-Kbp on the right side of *parS*_F_. Monte Carlo simulations and analytic formula are represented in red and dotted black lines, respectively. MC simulations are performed with a Freely-Jointed Chain of length N=4000 monomers of size *a*=10-bp, preventing finite size effect on the range of genomic coordinate considered. The cluster radius is *σ*=75nm and the binding energy difference *ε_ns_*-*ε_s_*=-0.9 is obtained by reading the density at the 1^st^ site after *parS* (see text and supplemental information). As a benchmark for simulations, the analytics are obtained from Eq.(1) with the same parameters. *Inset*; The ParB_F_ binding profile is represented as the number of nucleotide reads over 80-Kbp centered at *parS*. (D) and (E) Same as B and C with *parS*_F_ inserted at the *xylE* locus on *E. coli* chromosome from DLT3584 and DLT2075, respectively. The Kuhn length was adjusted to *a*=23-bp in the simulations and analytics. The characteristics of the A-F genetic loci are presented in Fig. S1A.

To study the assembly mechanism of partition complexes, we used the archetypical type I partition system of the plasmid F from *E. coli*. By varying several key parameters, we evaluated ParB binding patterns *in vivo* in relation to predictions of each model. We also investigated the chromosomal ParAB*S* system of the main chromosome of *Vibrio cholerae*. In all tested conditions, our data indicate that ParB binding profiles robustly correlate only with the predictions of the ‘Nucleation & caging’ model.

## Results

### ParB_F_ distribution pattern around *parS*_F_ is similar on chromosome and plasmid DNA

The plasmid F partition complex assembles on a centromere sequence, *parS*_F_, composed of twelve 43-bp tandem repeats (Helsberg and Eichenlaub, 1986), which contain ten 16-bp inverted repeat motifs to which ParB_F_ binds specifically *in vitro* (Pillet et al., 2011) and *in vivo* (Sanchez et al., 2015). Partition complex assembly has been investigated using small versions of the plasmid F, either ∼10- or ∼60-Kbp. To discriminate between the different partition complex assembly models, we used two larger DNA molecules: the native 100-Kbp plasmid F (F1-10B; Table S1) and the 4.6-Mbp *E. coli* chromosome with *parS*_F_ inserted at the *xylE* locus, in strains either expressing (DLT1472) or not (DLT1215) ParB_F_ from an IPTG-inducible promoter.

We first controlled the formation of ParB_F_ clusters on these two different DNA molecules using the ParB_F_-mVenus fluorescent fusion protein. ParB_F_-mVenus, fully functional in plasmid partitioning (Supplemental Table S2), was expressed from the endogenous locus on the plasmid F (F1-10B-BmV) or from a low-copy number plasmid under the control of an IPTG-inducible promoter (pJYB294). In both cases, we observed bright and compact foci in nearly all cells (Fig. 1B and D), indicating that the assembly of highly concentrated ParB_F_ clusters on *parS*_F_ from large DNA molecules, plasmid or chromosome, occurs similar to the smaller plasmid F counterparts (Sanchez et al., 2015). The number of foci from *parS*_F_ inserted on the chromosome is half of what is observed with the plasmid F, as expected from the twofold difference in copy-number (Collins and Pritchard, 1973).

We then performed ChIP-sequencing using anti-ParB antibodies and compared the ParB_F_ patterns from the 100-Kbp F1-10B plasmid and the *xylE*::*parS*_F_ chromosome insertion. For F1-10B, we observed a ParB binding pattern extending over 18-Kbp of *parS*_F_-flanking DNA nearly identical to the one previously observed on the 60-Kbp plasmid F (Sanchez et al., 2015), with the asymmetrical distribution arising from RepE nucleoprotein complexes formed on the left side of *parS*_F_ on *incC* and *ori2* iterons (Fig. 1C). When *parS_F_* is present on the chromosome, the ParB_F_ binding pattern displays a comparable enrichment of *xylE*::*parS*_F_flanking DNA over 15-Kbp (Fig. 1E). The ParB_F_ distribution extends ∼9- and 6-Kbp on the right and left sides of *parS*_F_, respectively. The asymmetry does not depend on *parS*_F_ orientation as an identical ParB_F_ binding pattern was observed with *parS*_F_ inserted in the reversed orientation (*xylE*::*parS*_F_-rev, Fig. S1B-C). On the left side, ParB_F_ binding ends near the *yjbE* locus that harbors two promoters (locus A; Fig. 1E, inset and S1A). On the right side, ParB_F_ binding ends at the *yjbI* gene locus (locus E; Fig. 1E and S1A). A dip in the ParB binding intensity is also observed ∼1-Kbp after *parS*_F_ spanning ∼300-bp, corresponding to a promoter region (locus C; Fig. 1E and S1A). Dips and peaks in this ParB_F_ binding pattern are different in terms of position and intensity when compared to the one present on the plasmid F. Overall, these data clearly indicate that the global ParB_F_ binding distribution around *parS*_F_ depends neither on the size nor the DNA molecule, plasmid or chromosome, and that the ParB_F_ binding probability is dependent on the local constraints of each given locus.

### The ‘Nucleation & caging’ binding model describes the partition complex assembly from the nucleation point to large genomic distance

Based on a smaller version of the plasmid F, we previously proposed the ‘Nucleation & caging’ model describing ParB stochastic binding at large distance (>100-bp) from *parS* due to DNA looping back into the confined ParB cluster. The characteristic asymptotic decay as a power-law with the exponent b=-3/2 is also observed with 100-Kbp plasmid F (Fig. 1C) and with *parS*_F_-inserted on the *E. coli* chromosome (Fig. 1E and Fig. S1C). This property is thus an intrinsic parameter of the ParB_F_ binding profile at distance >100-bp from *parS*_F_. The abrupt initial drop in ParB_F_ binding at a shorter genomic distance (<100-bp) from *parS*_F_ is explained by the difference of ParB_F_ binding affinities between specific *parS*_F_ sites (∼2 nM) and non-specific DNA (∼300 nM) (Ah-Seng et al., 2009). To take into account this initial drop, we now considered explicitly these different binding affinities: the amplitude of the drop, exp(ε_ns_ - ε_s_), is given by the ratio of the Boltzmann weights between specific (ε_s_) and non-specific (ε_ns_) binding energies (in units of *kT*). The ParB density was normalized to 1 by the value on the right side of *parS*, and captured in the following formula:

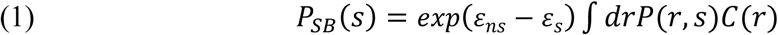

where 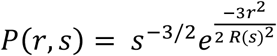 is the probability for two DNA loci spaced by a genomic distance *s* to be at a distance *r* in space for a Gaussian polymer; 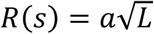 is the equilibrium size of DNA with linear length *L* and *α* is the Kuhn length of the DNA molecule (about twice the persistence length of the corresponding Worm-like chain; (Schiessel, 2013)); 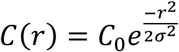 is the density of ParB at a radial distance *r* from the centromere, with *C*_0_ the concentration at the origin of the cluster and *σ* the typical size of the cluster. Note that *C*(*r*)*exp*(*ε_ns_* − *ε_s_*) is the linearized form of the Langmuir model (Phillips et al., 2012) offering a more compact and intuitive expression for *P(s)*. From (1) we easily calculate:

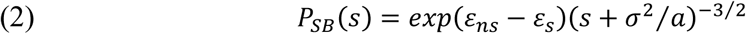

where the decay is asymptotically determined by a power law of exponent −3/2 modulated by an amplitude depending on the concentration and non-specific affinity of ParB. Two of the three parameters of this model are obtained from experiments: *σ*=75 nm is determined from superresolution microscopy (Lim et al., 2014; Sanchez et al., 2015) and *ε_ns_*-*ε_s_*=-0.9 is read directly from the ParB density at the nsDNA binding site after *parS* sequence. Note that *ε_ns_*-*ε_s_* estimate depends on the bioinformatics analysis (Fig. S1D). The only remaining free parameters is the Kuhn length *a*, set at 10- or 23-bp for the plasmid F or *parS*_F_-chromosomal insertions, respectively, to fully describe the ParB_F_ DNA binding profiles (Fig. 1C, E and Fig. S1C). These fitted values are lower than expected, likely due to the modeling that does not account for supercoiling. Nevertheless, using these defined parameters, the refined ‘Nucleation & caging’ model provides a qualitative prediction of the experimental data over the whole range of genomic positions, from a few bp to more than 10-Kbp.

### ParB_F_ DNA binding pattern over a wide range of ParB concentrations favors the ‘Nucleation & caging’ model

The physical modeling for each proposed model (Broedersz et al., 2014; Sanchez et al., 2015) predicts distinct and characteristic responses upon variation of the intracellular ParB concentration (see explanations in Fig. S2A). Briefly, (i) the ‘1-D filament’ model predicts a rapid decrease of ParB binding followed by a constant binding profile dependent on ParB amount, (ii) the ‘Spreading & bridging’ model predicts linear decays with slopes depending on the ParB amount, and (iii) the ‘Nucleation & caging’ model predicts a binding profile which depends only on the size of the foci. The exponent b=-3/2 of the power-law distribution would not change upon ParB amount variation resulting in an overall similar decay. In order to discriminate between these three model predictions, we performed ChIP-seq experiments over a large range of intracellular ParB concentrations. To prevent interference with plasmid stability, we used the chromosomally encoded *xylE*::*parS*_F_ construct expressing *parB*_F_ under the control of an IPTG inducible promoter (DLT2075).

Without IPTG induction, ParB_F_ was expressed at ∼0.2 of the physiological concentration from plasmid F, as judged by Western blot analyses (Fig. S2B). We also tested an 8- and 14- fold overproduction of ParB_F_. Assuming the two-fold difference in copy number (Fig. 1B and 1D), these three conditions provided ParB_F_/*parS*_F_ ratios of 0.4, 16 and 28, relative to the plasmid F one. At these three ratios, ChIP-seq data revealed that ParB_F_ binding extended similarly over ∼15-Kbp around *parS*_F_. We analyzed the right side of *parS*_F_ displaying the longest propagation distance by normalizing each data set (Fig. 2A). It revealed that regardless of ParB concentration (i) the ParB distribution in the vicinity of *parS*_F_ always displays a good correlation with a power law fitting with an exponent of −3/2, (ii) the ParB binding profile ends at the same genomic location, i.e. 9-kpb from *parS*_F_ and (iii) the dips and peaks in the pattern are highly conserved. This indicates a highly robust ParB binding pattern that is invariant over a ∼70-fold variation of the ParB amount.

To further vary the amount of ParB_F_ available for partition complex assembly, high-copy number (HCN) plasmids containing the *parS*_F_ sequence were introduced into the *xylE*::*parS*_F_ strain to efficiently titrate ParB_F_ by its binding to the excess of specific binding sites (∼200- and ∼500-fold on pBR322 and pBSKS derivatives, respectively; (Diaz et al., 2015)). Epifluorescence microscopy of these strains reveals that all cells display a diffuse ParB-mVenus fluorescence (Fig. 2B) in contrast to concise foci without titration (Fig. 1A), suggesting a large reduction of ParB availability to non-specific sites in the vicinity of *parS*_F_ on the chromosome. ChIP-seq analyses in the two titration conditions revealed that ParB binding in the vicinity of *parS*_F_ was dramatically reduced as expected. However, rescaling the signals by a factor of 10 and 50 for the pBR322 and pBSKS *parS*_F_-carrying derivatives, corresponding to a ParB_F_/*parS*_F_ ratio of 0.04 and 0.016, respectively, revealed a ParB_F_ binding pattern above the background level (Fig. 2B, inset). In both datasets, ParB_F_ binding decreases progressively over about the same genomic distance and with a similar power law decay as without titration. Moreover, even with these very low amounts of available ParB_F_, the dips and peaks in the profiles are present at the same positions.

**Figure 2.**
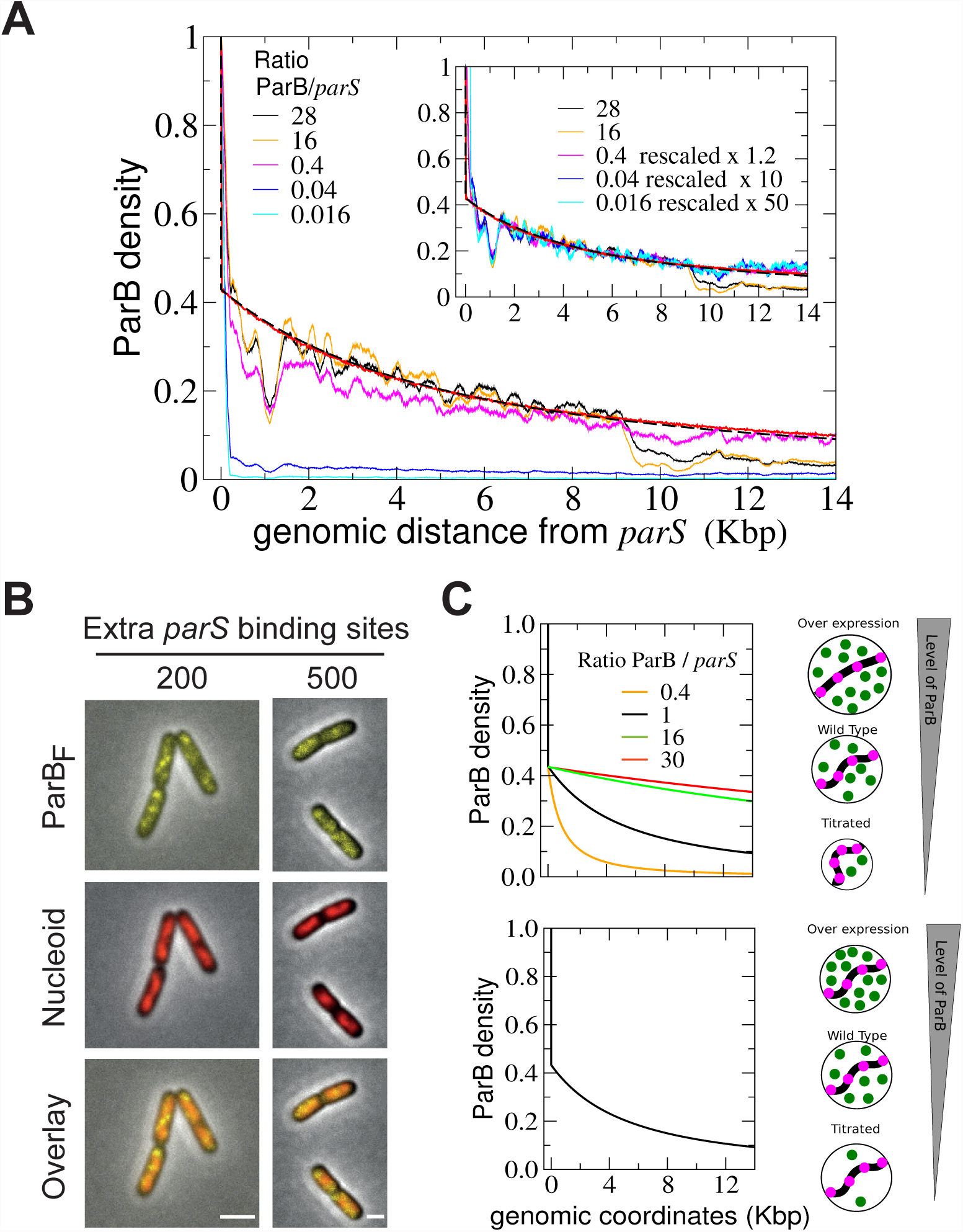
ParB_F_ DNA binding pattern is robust over a large range of intracellular ParB_F_ concentrations. (A) Normalized ParB_F_ binding profiles at different ParB_F_/*parS*_F_ ratio. ChIP-seq density on the right side of *parS*_F_ inserted at *xylE* were measured in DLT2075 induced (16, 28) or not (0.4) with IPTG, or carrying HCN plasmids pZC302 (0.04) or pJYB57 (0.016). The ParB/*parS* ratio is calculated relative to the one of plasmid F as determined from Western blot analyses (Fig. S2B). Monte Carlo simulations and analytical formula are plotted with the same parameters as in Fig. 1E. *Inset*; the amplitudes of the curves are rescaled by the indicated factors to overlap with the curves of highest amplitude. (B) ParB_F_ are dispersed in the cell upon titration by HCN plasmids. ParB_F_-mVenus expressed from pJYB294 were imaged as in Fig. 1D in DLT3577 (left) and DLT3576 (right) carrying pZC302 and pJYB57, respectively. The number of extra *parS*_F_ per cell, indicated on top of each raw, are estimated from the copy number per cell of HCN plasmids carrying 10 specific binding sites. White bars: 1 µm. (C) The size of ParB_F_ clusters is independent of the intracellular ParB_F_ concentration. We considered two possible evolutions of the cluster size upon variations of ParB amount in the framework of ‘Nucleation & caging’ with corresponding schematics drawn on the right. *Top*: constant ParB concentration; supposing that clusters are compact, the cluster radius *σ* would depend on the number *m* of ParB like *σ* = m^1/3^. Predictions profiles, plotted at different ratio of ParB/*parS*, vary within the range of the experimental levels tested. *Bottom*: constant cluster size; ParB concentrations vary but the range of exploration remains the same resulting in overlapping prediction profiles.

The invariance of the overall ParB profile over three orders of magnitude of ParB concentration (Fig. 2B, inset) excludes the predictions of both the ‘1-D filament’ and the ‘Spreading & bridging’ models (Fig. S2A). In addition, the conservation in the positions of the dips and peaks indicate that the probability of ParB_F_ binding at a given location is also not dependent on the amount of ParB_F_ in the clusters. These results are strongly in favor of the refined ‘Nucleation & caging’ model presented above.

### The size of the dynamic ParB/*parS* cluster is independent of ParB intracellular concentration

In all of the ParB induction levels tested, the genomic distance over which ParB_F_ binds around *parS*_F_ is constant and displays a very similar decay (Fig. 2A). This conserved binding behavior could provide information on the cluster size as a function of ParB amount. Indeed, the ‘Nucleation & caging’ model predicts a probability *P*(*s*) ∼ (*s*+*C*) ^-3/2^ of ParB binding at a genomic distance *s*, where the constant *C* = *σ*^2^/*a* is function of the average radius of the foci *σ* and the Kuhn length of the DNA *a*. Thus, the *P*(*s*) decay is entirely determined by the geometry of the foci and the intrinsic flexibility of the DNA. Varying the ParB amount could lead to two situations: (i) the density of ParB, but not *σ*, is constant (ii) *σ* is fixed and ParB density is variable. We plotted these two situations in the range of ParB/*parS* ratio considered experimentally (Fig. 2C): with (i), the different *P(s)* strongly varied, and (ii), *P(s)* was invariant relative to the ParB amount resulting in overlapping profiles. Experimental data (Fig. 2A) are in excellent agreement with the latter. From this modeling, we thus concluded that the size of partition complexes is invariant to change in ParB intracellular concentration.

### The arginine rich motif (box II) of ParB_F_ is critical for partition complex assembly

The ability of ParB to multimerize through dimer-dimer interactions is required for the formation of ParB clusters. A highly-conserved patch of arginine residues present in the N-terminal domain of ParB (box II motif; Yamaichi and Niki, 2000) has been proposed to be involved in ParB multimerization (Breier and Grossman, 2007; Song et al., 2017). To examine to what extent the box II motif is involved *in vivo* in the assembly of ParB_F_ clusters, we changed three arginine residues to alanine (Fig. S3A). The resulting ParB_F_-3R* variant was purified and assayed for DNA binding activity by electro-mobility shift assay (EMSA) in the presence of competitor DNA using a DNA probe containing a single *parS*_F_ site (Fig. 3A). ParB_F_-3R* binds *parS*_F_ with high affinity (B1 complex) indicating no defect in (i) protein folding, (ii) *parS*_F_ binding and (iii) dimerization, a property required for *parS* binding (Hanai et al., 1996). However, in contrast to WT ParB, the formation of secondary complexes (B’2 and B’3), proposed to result from ParB multimerization (Sanchez et al., 2015), was impaired further suggesting the implication of box II in dimer-dimer interaction. A mini-F carrying the *ParB_F_-3R\** allele (pAS30) was lost at a rate corresponding to random distribution at cell division (Table S2), indicating that this variant is unable to properly segregate the mini-F.

The ParB_F_-3R* variant was then expressed in native or fluorescently-tagged (ParB -R3*- mVenus) forms, from pJYB303 or pJYB296, respectively, in the *xylE*::*parS*_F_ strain. By imaging ParB_F_-3R*-mVenus, we observed only faint foci in a high background of diffuse fluorescence (Fig. 3B). These barely detectable foci may correspond to ParB_F_-3R*-mVenus binding to the ten specific sites present on *parS*_F_ and, if any, to residual ParB_F_ cluster formation. We then performed ChIP-seq assays with ParB_F_-3R* present in ∼25-fold excess (relative ParB_F_/*parS*_F_ ratio compared to the plasmid F one; Fig. S3B). The resulting DNA binding profile displayed enrichment only at *parS*_F_ with a total absence of ParB_F_ binding on *parS*_F_-flanking DNA (Fig. 3C). This pattern differs from those observed in conditions of ParB_F_ titration (Fig. 2A; inset), indicating that the ParB_F_-3R* box II variant is fully deficient in clustering *in vivo*. The same pattern was also observed with ParB_F_-3R*-mVenus (Fig. S3C) indicating that the mVenus fluorescent-tag fused to ParB_F_ does not promote cluster assembly.

**Figure 3.**
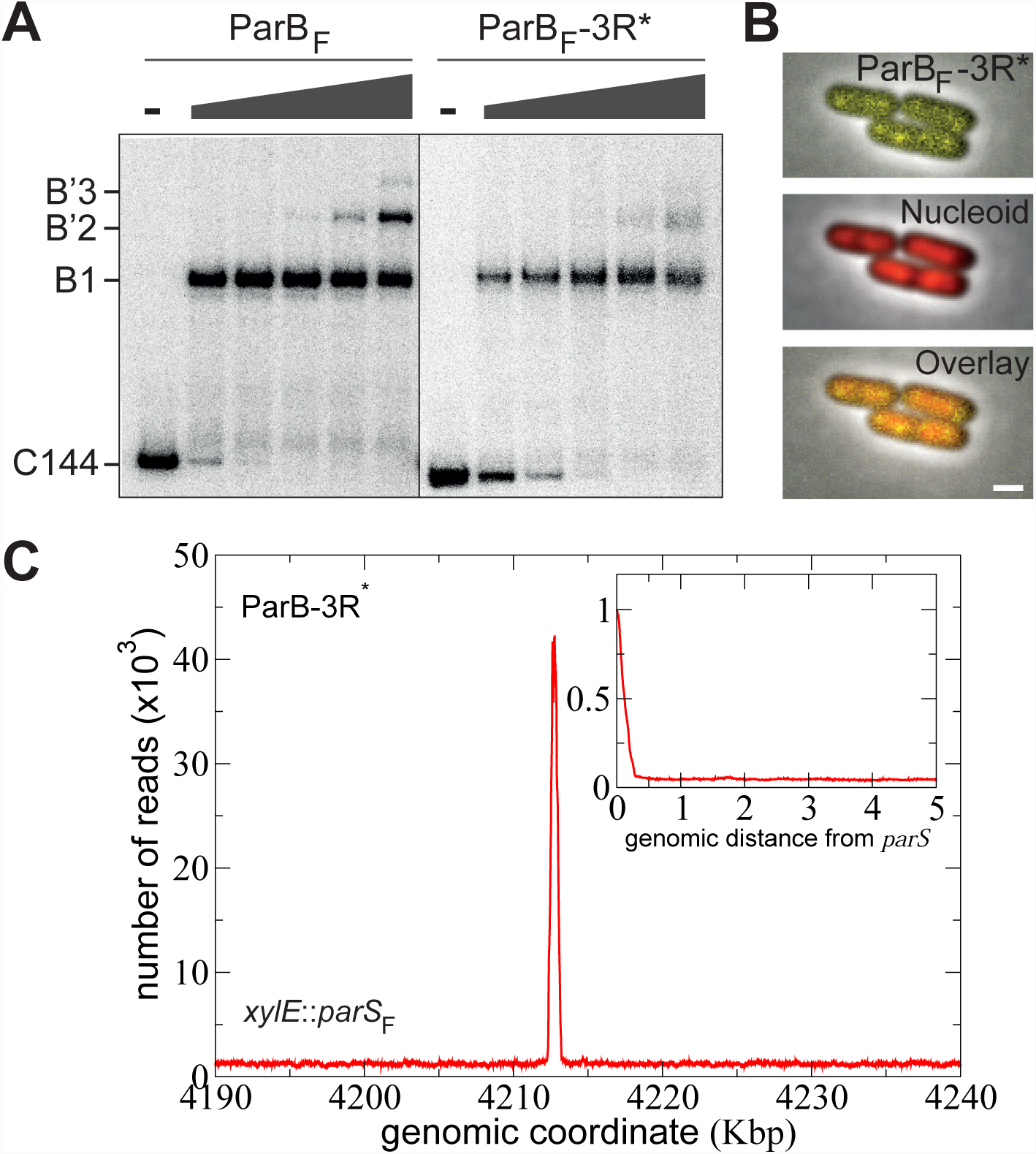
The box II motif of ParB_F_ is crucial for ParB_F_ binding in the vicinity of *parS*_F_ and cluster formation. (A) The formation of secondary ParB_F_-DNA complexes requires the box II motif. EMSA were performed with a 144-bp ^32^P-labelled DNA fragments (C144) carrying a single 16-bp *parS* binding motif. Reaction mixtures containing 100 µg.ml^−1^ sonicated salmon sperm DNA were incubated in the absence (-) or the presence of increasing concentrations (grey triangle; 10, 30, 100, 300, and 1000 nM) of ParB_F_ or ParB_F_-3R*. Positions of free and bound probes are indicated on the left. B1 represents complexes involving the specific interaction on the 16- bp binding site, while B’2 and B’3 complexes represent secondary complexes involving the *parS*_F_ site with one or two additional nsDNA-binding interactions, respectively. (B) ParB_F_ cluster formation requires the box II motif. Epifluorescence microscopy of ParB_F_-3R*-mVenus from DLT3566 is displayed as in Fig. 1D. White bars: 1 µm. (C) ParB_F_ *in vivo* DNA binding in the vicinity of *parS*_F_ sites requires the box II motif. ChIPseq was performed on DLT3726 carrying *parS*_F_ in the *xylE* chromosomal locus and expressing ParB_F_-3R* variant. ParB_F_-3R* DNA binding profile displayed the number of nucleotide reads as a function of the *E. coli* genomic coordinates. The peak at *parS*_F_ covered approximately 950-bp, which corresponds to the 402-bp between the 1^st^ and 10^th^ specific binding sites and ∼280-bp on each sides (representing the average size of the DNA library; see Fig. S3D). No ParB_F_-3R* enrichment was found on *parS*F-flanking DNA and elsewhere on the chromosome. *Inset*, zoom in on the right side of *parS*_F_ over 5-Kbp with the ParB density, normalized to 1 at the first bp after the last *parS* binding repeat, plotted as a function of the distance from *parS*_F_.

Together, these results indicate that the box II variant is specifically deficient in ParB_F_ cluster assembly but not in *parS*_F_ binding, and thus reveal that the box II motif is critical for the auto-assembly of the partition complex.

### ParB also propagates stochastically from native chromosomal *parS* sites

ParABS systems are present on most bacterial chromosomes (Gerdes et al., 2000). To determine whether chromosomal ParB-*parS* partition complexes also assembled *in vivo* in a similar manner to the plasmid F, we investigated the bacterium *Vibrio cholerae*, whose genome is composed of two chromosomes. We focused on the largest chromosome to which ParB*_Vc_*_1_ binds to three separated 16-bp *parS* sites comprised within 7-Kbp (Baek et al., 2014; Saint-Dic et al., 2006) (Fig. 4A).

We purified ParB*_Vc_*_1_ antibodies against his-tagged ParB*_Vc_*_1_ and performed ChIP-seq assays on exponentially growing cultures. The ParB*_Vc_*_1_ DNA binding pattern covered ∼18- Kbp and displayed three peaks at the exact location of the three *parS_Vc_*_1_ sites (Fig. 4B). Each peak exhibits a distinct but reproducible difference in intensity that might correspond to the slight differences in *parS_Vc_*_1_ sequences (Fig. S4A). An asymmetry in the binding pattern was observed on the left side of *parS*1 with the limit of ParB*_Vc_*_1_ binding corresponding to the end of the rRNA operon located ∼4-Kbp upstream from *parS*1 (Fig. 4B). This suggests that highly transcribed genes might significantly interfere with the extent of ParB binding.

**Figure 4.**
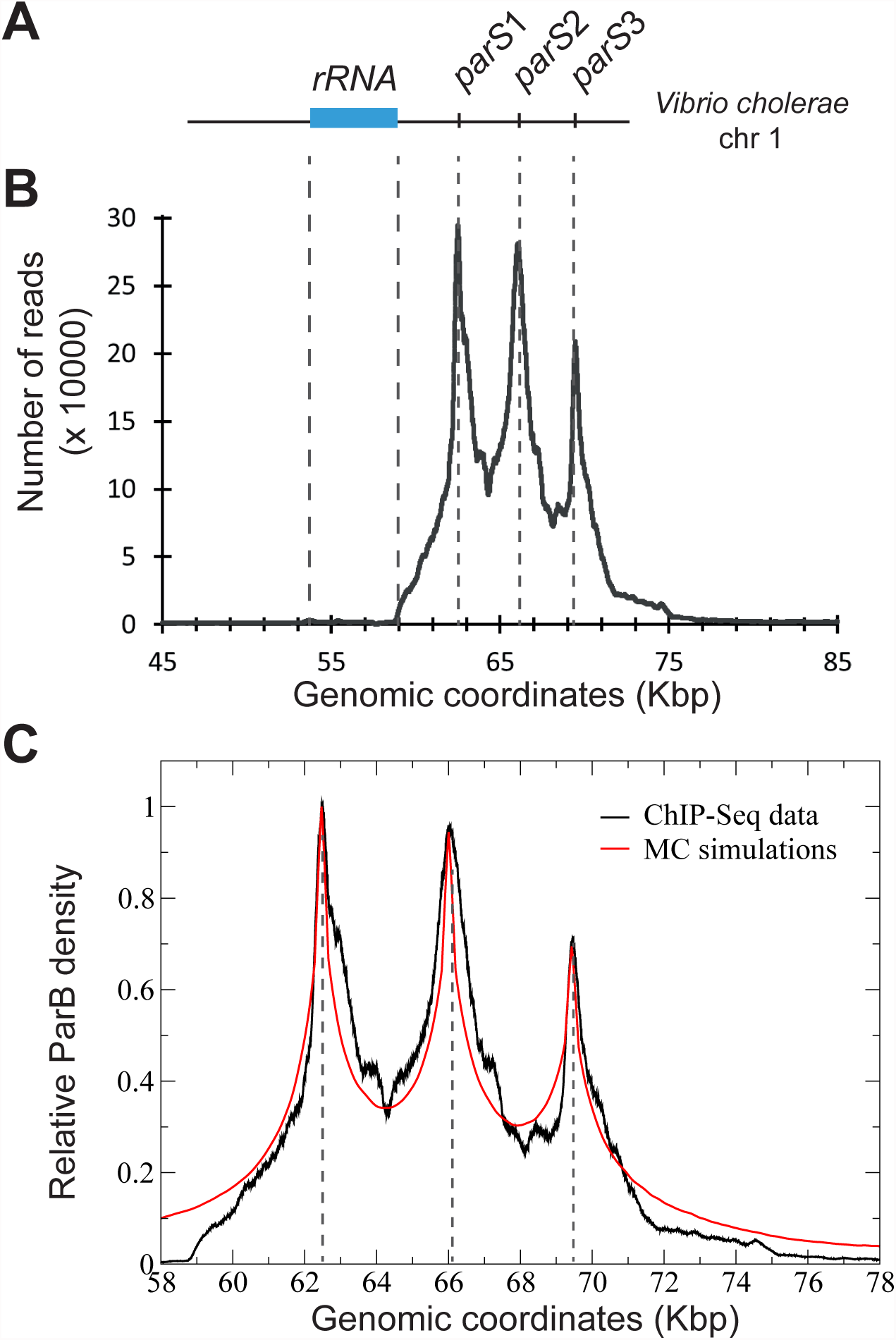
ParB of *V. cholerae* assembled in cluster similarly to ParB_F_. (A) Schematic representation of the genomic locus of the chromosome 1 of *V. cholerae* with the three *parS* sites, named *parS*1 -3. The rRNA operon (blue rectangle) spans the genomic coordinates 53823 to 59123. (B) ChIP-seq performed on strain N16961 is displayed as the number of nucleotide reads in function of the genomic coordinates. Correspondence to the *parS*1 -3 location in (A) is indicated by grey dotted lines. (C) We model the ChIP-Seq data as in Fig. 1C-E by means of MC simulations with a Freely jointed chain of size *N*=2000 monomers of size *a*=16-bp. The best fit was achieved with *σ*=25nm and the difference *ε_ns_*-*ε_s1_*=-0.2 between non-specific binding energy and chemical potential (read directly from ChIP-Seq data). In the MC simulation, we accounted for the finite width of the distribution around *parS* sites by including the average size of the DNA library (304-bp; Fig. S1D and S4B).

We modeled ParB*_Vc_*_1_ DNA binding profile with the framework of the refined Stochastic Binding model (see above), considering three non-interacting spheres centered on each of the *parS* sites (Fig. 4C). Here, *ε_ns_*-*ε_s1_*=-0.2, where *ε_s1_* is the specific binding energy for *parS1*. The simulated profile was obtained by using the same protocol as performed by the bioinformatic analysis in order to account for the width of the peaks around each *parS*; the same modeling as for *E. coli* would led to a sharp decay between *parS* and non-specific sites (Fig. S4B).

The maxima in the ParB binding profile depends on the *parS* sites (Fig. 4C) and are interpreted as a difference in binding affinity. In the simulations, the ParB density is normalized to 1 by the value on the right of *parS*1. The relative density of the two other p*arS* sites is fixed according to the values read on the ChIP-seq plot (3% and 29% lower affinity for *parS*2 and *parS*3 compared to *parS*1, respectively). We found a good agreement with the ParB*_Vc_*_1_ profile by applying a lower difference between the specific and non-specific binding energies than for ParB_F_, as reported in other ParAB*S* system (Fisher et al., 2017). We also noticed a clear difference at the minima of ParB binding on either side of *parS*_2_ (64.2 and 68- Kbp; Fig. S4B). In the case of a single cluster constraining the three *parS*, the profile would only depend on the genomic distance from *parS*2 resulting in a symmetrical pattern, while in the case of three independent clusters, an absence of symmetry due to the occupation of the specific sites is expected. This indicates that the system displays three independent clusters nucleated at each *parS* sites. However, the possibility that these clusters mix together at a frequency dependent on the genomic distance between *parS* sites is not excluded. At larger distances from *parS* sites, differences between the experimental data and the simulation probably arise from strong impediments to ParB binding, such as the presence of the rRNA operon.

These data strongly support that the partition complex assembly mechanism is conserved on plasmid and chromosome ParAB*S* systems.

### Nucleoprotein complexes, but not active transcription, are the major determinants for the impediment of ParB stochastic binding

The major dips in the ParB_F_ DNA binding signal are often found at promoter loci (Fig. S1A). To investigate the link between gene expression and the impediment to ParB propagation, we reproduced the ChIP-seq assays using the *xylE*::*parS*_F_ strain grown in the presence of rifampicin, an inhibitor of RNA synthesis that traps RNA polymerases at promoters loci in an abortive complex unable to extend RNAs beyond a few nucleotides (Herring et al., 2005). We did not observe significant changes to the ParB signal on either side of *parS*_F_ (Fig. 5A; compare red and blue curves). Notably, the ParB signal still strongly drops in promoter regions (e.g. loci A, C and E) and the dips and peaks are present at the same locations (Fig. 5B). This indicates that active transcription by RNA polymerase is not a major impediment to ParB binding.

**Figure 5.**
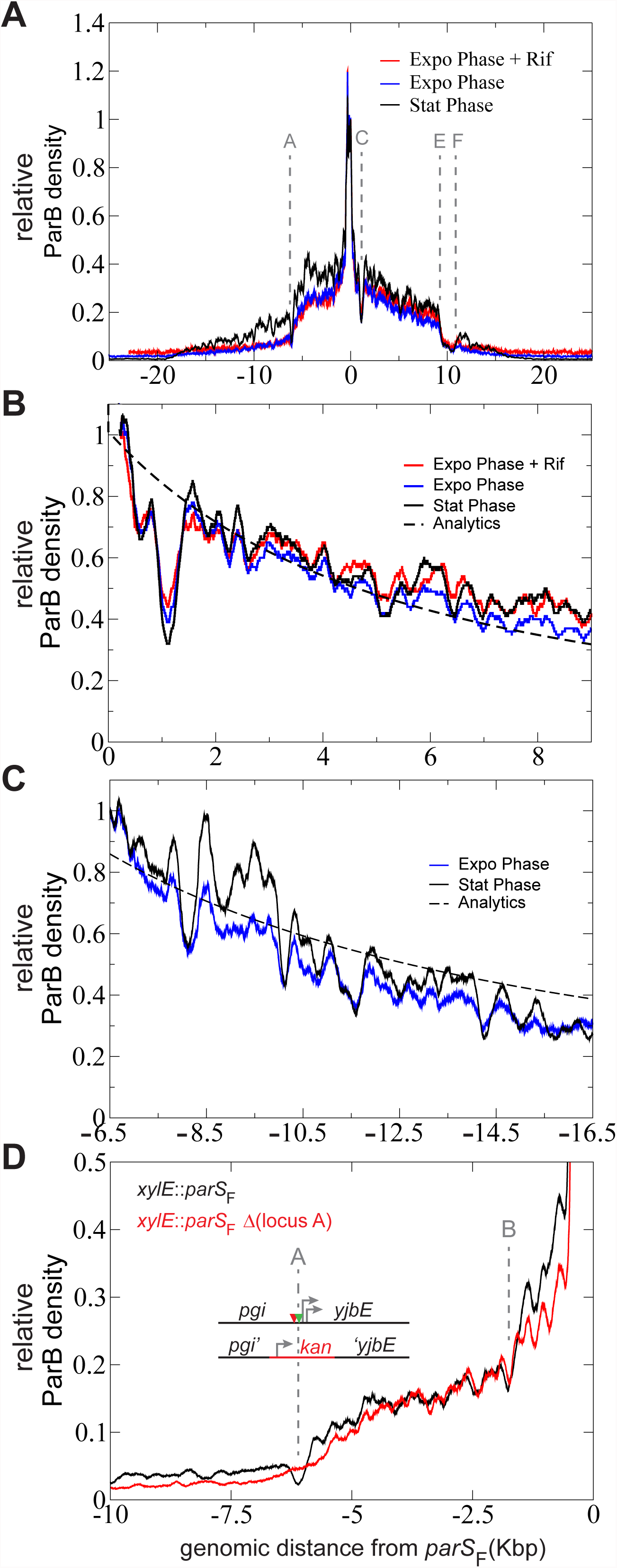
Robust dips and peaks signatures in ParB DNA binding profiles. ChIP-sequencing assays were performed on DLT2075 (*xylE*::*parS*_F_) expressing ParB_F_ grown in exponential (expo) or stationary (stat) phases with addition of rifampicin when indicated (+Rif). (A) ParB_F_ DNA binding around *parS*_F_ is independent of active transcription. The color-coded ParB_F_ profiles are represented over 50-Kbp as the relative ParB density normalized to 1 at the first bp after the last *parS*_F_ binding site. Loci A, C, E and F are defined in Fig. S1A. (B) The dips and peaks are highly similar in the three indicated conditions. Same as in (A) with zoom in on the right side of *parS*_F_ up to 9-Kpb and normalization to 1 at genomic coordinate 230. The dotted line corresponds to the analytics description of Stochastic Binding (see details in Fig. 1C-E). (C) ParB_F_ binding profile upstream of the locus A. Same as in (A) with zoom in from −6.5 to - 16.5-Kbp and normalization to 1 at genomic coordinate −6.5-Kbp (upstream of the dip at the locus A). ParB_F_ binding profile follows the power-law characteristic of stochastic binding, represented by the analytics description (dotted line), upstream of the locus A in stationary phase (black) and in exponential phase (blue). (D) The promoter region at locus A prevents ParB_F_ DNA binding. Chip-seq assays were performed in isogenic *xylE*::*parS*_F_ strains (DLT2075; black curve) in which the locus A is replaced by a kanamycin gene (DLT3651; red curve). The relative ParB density as a function of the distance from *parS*_F_ is drawn and normalized as in (A). The promoter region is depicted as in Fig. S1A.

We also measured the ParB binding profile in stationary phase, a growth condition in which gene expression is strongly reduced. On the right side of *parS*_F_, ParB distribution was similar to all other tested conditions (Fig. 5A), thus confirming the robustness of the binding pattern. On both sides, the strong reduction of ParB binding at loci A, C and E was still observed. However, in contrast to the other conditions, ParB binding recovers after these loci and extends up to ∼18-Kbp on both sides, resulting in the location of *parS*_F_ in the middle of a ∼36-Kbp propagation zone. Interestingly, the ParB binding profiles after these recoveries could still be fit to a power law exhibiting the same characteristics as at lower genomic distances (Fig. 5C). In stationary phase, the reduced intracellular dynamics (Parry et al., 2014) and the higher compaction of the DNA (Meyer and Grainger, 2013) may stabilize the partition complex revealing the ParB_F_ bound at larger distances from *parS*_F_. Interestingly, in higher (stationary phase) or lower (rifampicin-treated cells) DNA compaction states (Fig. S5A), the ParB_F_ DNA binding pattern is not altered, exhibiting a similar profile of dips and peaks (Fig. 5B). This indicates that the assembly of the partition complex is not perturbed by variation in DNA compaction level within the nucleoid.

To further demonstrate the impediment of ParB_F_ binding in promoter regions, we constructed a strain in which the locus A, carrying two promoters, an IHF and two RcsB binding sites, is replaced by a kanamycin resistance gene (Fig. 5D). The measured ParB_F_ binding pattern remained highly comparable except at the locus A where the dip is absent. This result clearly indicates that site-specific DNA binding proteins are the main factors for restricting locally ParB_F_ binding.

### ParB molecules exchange rapidly between partition complexes

Single molecule *in vivo* localization experiment have shown that over 90% of ParB_F_ molecules are present at any time in the confined clusters (Sanchez et al., 2015), suggesting that partition complexes are stable structures. However, stochastic binding of most ParB_F_ on non-specific DNA suggests that partition complexes are highly dynamic. To reconcile this apparent discrepancy, we performed fluorescence recovery after photobleaching (FRAP) on two foci cells for measuring ParB_F_ dynamics between partition complexes. By laser-bleaching only one focus, we could determine whether ParB_F_ dimers could exchange between clusters and measure the exchange kinetics. As ParB_F_ foci are mobile, we choose to partially bleach (∼50%) the focus enabling immediate measurement of fluorescence recovery (Fig. 6A-B). A few seconds after bleaching, the fluorescence intensity recovers while it decreases in the unbleached focus. This exchange is progressive and the intensity between the two foci equilibrated in ∼80 sec on average (between 50 and 120 sec for most individual experiments). We estimate that, when exiting a cluster, each ParB_F_ dimer has the same probability to reach any of the two clusters. Therefore, the time of equilibration between the two foci corresponds to the exchange of all ParB_F_. These results thus indicate that the partition complexes are dynamic structures with a rapid exchange of ParB_F_ molecules between clusters.

**Figure 6.**
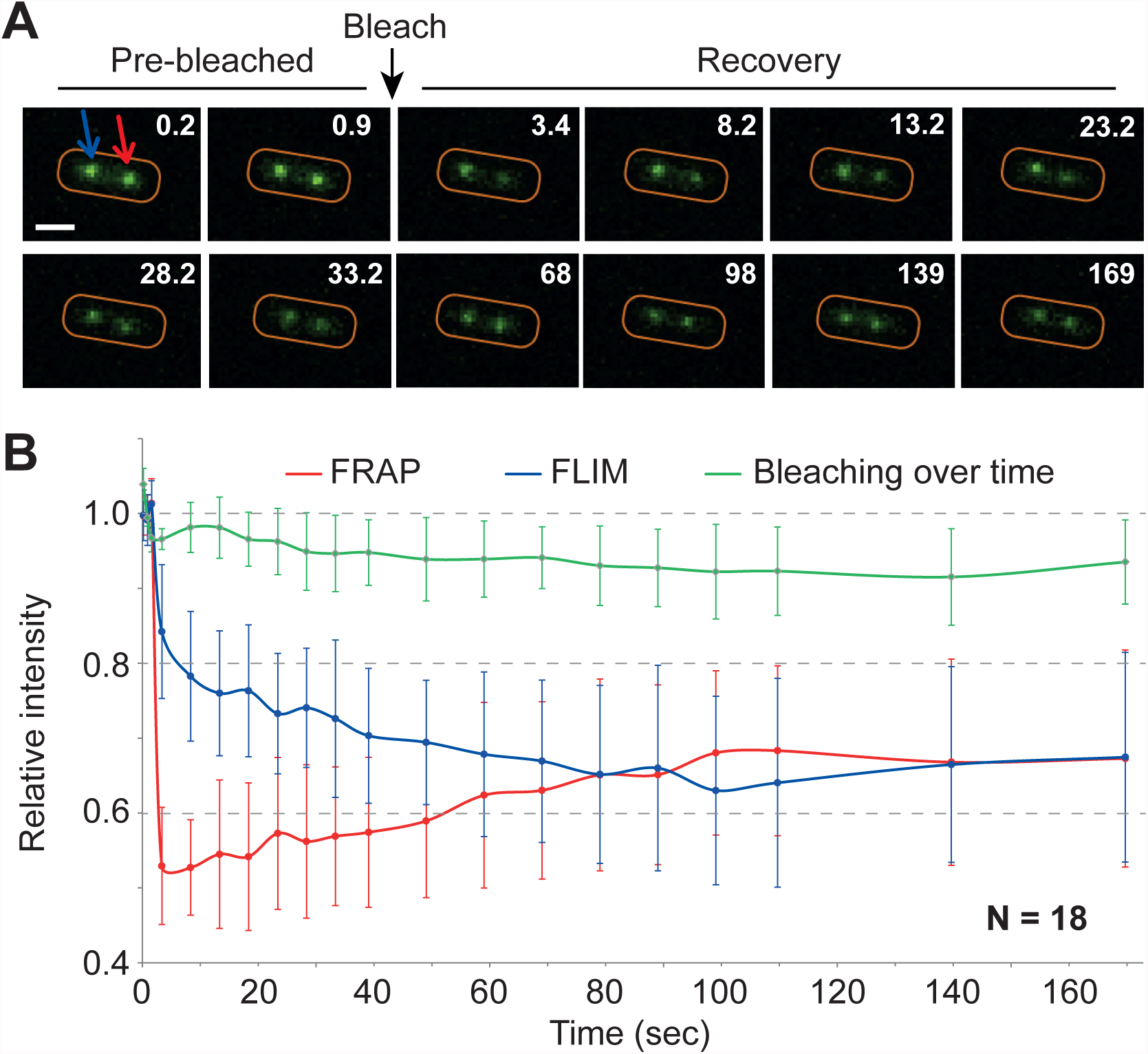
ParB dynamics between partition complexes. ParB_F_ exchange between foci was measured by FRAP and FLIM (fluorescence lifetime imaging microscopy) from two-foci cell of DLT1215 carrying pJYB234. A- Representative images of a photobleached cell during a FRAP experiment. The 488 nm laser was pulsed (Bleach) on one of the two foci at ∼2.4 sec (black arrow). Red and blue arrows correspond to the bleached and unbleached focus, respectively. Time is indicated in second (upper right). The cell outline is drawn in red. Scale bar: 1 μm. B- Quantification of ParB_F_-mVenus fluorescence intensity over time. The dynamics of fluorescence intensity is shown from averaging 18 independent measurements of the bleached (FRAP, red line) and unbleached (FLIM, blue line) foci. Foci fluorescence intensity in each experiment was normalized to the average intensity of each focus before photobleaching. Natural bleaching during the course of the experiments (green curve) was estimated for each measurement by averaging the fluorescence intensity of 15 foci present in each field of view.

## Discussion

Despite over three decades of biochemical and molecular studies on several ParAB*S* systems, the mechanism of how a few ParB bound to *parS* sites can attract hundreds of ParB in the vicinity of *parS* to assemble a high-molecular weight complex remained puzzling. The three main mechanisms proposed for ParB-*parS* cluster formation have been studied from physico-mathematical perspectives (Broedersz et al., 2014; Sanchez et al., 2015), predicting very different outcomes for the ParB binding profile in the vicinity of *parS* sites upon change in ParB concentration. Here, the ParB binding patterns were found invariant over a large variation of ParB amount displaying a robust decay function as a power law with the characteristic exponent b=-3/2 and a conserved length of the propagation zone (Fig. 2A). Strikingly, even in the titration conditions tested, which resulted in a very low amount of ParB available to bind to non-specific DNA sites, the overall ParB DNA binding pattern remained invariant (Fig. 2A, inset). Neither ‘1-D spreading’ nor ‘Spreading & bridging’ physical models could describe these data in the conditions tested (Broedersz et al., 2014). A variant of the latter model has explored the ParB binding pattern in the low spreading strength limit (Walter et al., 2018). This ‘Looping & clustering’ model also predicts variations in the ParB binding pattern over a simulated 4-fold range of ParB amount, which is in contrast to the invariant pattern observed experimentally over more than three orders of magnitude (Fig. 2). In conclusion, only the ‘Nucleation & caging’ model based on stochastic ParB binding well describes the experimental data and provides accurate predictions for the mechanism of the partition complexes assembly.

We refined the modeling of the dynamic and stochastic ParB binding model by including DNA binding affinities for specific and non-specific sites to describe the initial drop observed immediately after *parS*. In this framework, we found that ParB clusters have a constant size accommodating important variations in ParB concentration (Fig. 2C). We propose that the cluster size is dependent on the intrinsic ParB-ParB and ParB-nsDNA interactions, and would thus be an inherent characteristic of each ParAB*S* system (Funnell and Gagnier, 1993; Sanchez et al., 2015; Taylor et al., 2015). The refined modeling also well describes the chromosomal partition system of *V. cholera*, predicting three independent clusters nucleated at each of the three *parS* sites (Fig. 4C). In all cases reported here, the partition complex assembly is well described by the ‘Nucleation & caging’ model, and we propose that this mechanism of assembly is conserved on chromosome and plasmid partitioning systems.

In addition to its robustness within a large range of ParB concentration (Fig. 2A) and different nucleoid compaction states (Fig. 5A), the *in vivo* ParB DNA binding pattern also exhibits conserved dips and peaks at particular locations. The major dips are located at promoter regions (Fig. 1E and S1A) but do not depend on active transcription (Fig. 5B). This suggests that these specific signatures mostly depend on the intrinsic local genomic environment. This hypothesis was confirmed by deleting the locus A, carrying several regulator binding sites, which led to the suppression of the dip at this position (Fig. 5D). Therefore, proteins such as transcriptional regulators and NAPs (nucleoid associated proteins) that bind specifically to DNA prevent ParB binding to these sites, thus reducing locally the ParB signal. We propose that this impediment to ParB binding is proportional to the time of occupancy of these regulators at their site-specific DNA binding sites. Larger nucleoprotein complexes, as exemplified on the plasmid F at the iteron sites (*ori2* and *incC*; Fig. 1C) that interact *in cis* and *in trans* (Das and Chattoraj, 2004), were previously proposed to be spatially excluded from the vicinity of the ParB cluster with a low probability that DNA beyond these sites comes back into the cluster preventing ParB binding (Sanchez et al., 2015). Such an exclusion does not occur from smaller protein-DNA complexes, with the recovery of the ParB binding signal that further follows the characteristic power law decay (e.g. locus A; Fig. 5C). These results show that low molecular weight protein-DNA complexes do not impair the overall, only the local, ParB binding pattern.

The formation of highly concentrated clusters of ParB relies on a strong ParB-*parS* interaction and two other interactions, ParB-ParB and ParB-nsDNA (Fisher et al., 2017; Sanchez et al., 2015). ParB mutants that do not propagate outside *parS* are impaired in partition activity and in cluster formation *in vivo* (Breier and Grossman, 2007; Rodionov et al., 1999). The conserved box II motif (Yamaichi and Niki, 2000) was suggested to be part of the dimer-dimer interface (Breier and Grossman, 2007; Graham et al., 2014) but some misfolding caveat has been reported with some mutants, such as ParB*_Bsub_*_-G77S_ (Song et al., 2017). *In vivo* the box II variant (ParB_F_-3R*) is totally deficient in partition activity and cluster formation (Fig. 3B) while proficient for *parS*_F_ binding (Fig. 3C). The total absence of ParB_F_-3R* binding outside *parS*_F_ (Fig. 3C and S3D) indicates that the box II motif is the major interface for the interaction between ParB dimers and is critical for the partition complexes assembly *in vivo* and the DNA partition activity.

ParA interacts with partition complexes in a ParB-dependent manner both *in vitro* and *in vivo* (Bouet and Funnell, 1999; Lemonnier et al., 2000) to ensure the ATP-dependent segregation of centromere sites upon DNA replication (Ah-Seng et al., 2013; Fung et al., 2001; Scholefield et al., 2011). Previous studies from *V. cholerae* and *S. Venezuela* have reported contradictory results on the involvement of ParA in the assembly of the partition complex (Baek et al., 2014; Donczew et al., 2016), which may arise from the pleiotropic effects of ParA on cellular processes, such as gene transcription or DNA replication (Murray and Errington, 2008). The ParB_F_ DNA binding profiles on the plasmid F (Fig. 1C) and on the *E. coli* chromosome (Fig. 1E), in the presence and absence of ParA_F_, respectively, are highly similar, therefore indicating that they assemble independently of ParA. Partition complexes, composed of hundreds of ParB dimers, were thought to be confined at the interface between the nucleoid and the inner membrane (Vecchiarelli et al., 2012). The observation that they rather are located within the nucleoid in a ParA-dependent manner (Le Gall et al., 2016) raises the question as to how they are not excluded from it. The ‘Nucleation & caging’ model could solve this apparent paradox. Indeed, relying on a strong ParB-*parS* interaction (nM range) and two other synergistic, but labile interactions, ParB-ParB and ParB-nsDNA (hundreds of nM range; Fisher et al., 2017; Sanchez et al., 2015), it would allow the dynamic confinement of most ParB without forming a rigid static structure. This dynamic organization is further supported by the finding that ParB dimers quickly exchange between clusters (∼80 sec; Fig. 6). By comparison, the equilibration times between H-NS or TetR-*tetO* clusters were 5 or 10 times much longer, respectively (Kumar et al., 2010). Since >90% of ParB are present in clusters (Sanchez et al., 2015), it implies that their time of residency is much longer inside than outside, in agreement with fast diffusion coefficients (∼ 1 µm^2^. s^−1^) for non-specific DNA binding proteins (Kumar et al., 2010). We propose that, collectively, all the individual but labile interactions for partition complex assembly allow the whole complex attracted by ParA to progress within the mesh of the nucleoid.

## Experimental procedures

### Bacterial strains and plasmids

*E. coli* and *V. cholerae* strains and plasmids are listed in Supplemental Table S1. Plasmids and strains constructions, growth cultures and plasmids stability assays are described in Supplemental experimental procedures.

### Epifluorescence microscopy

Exponentially growing cultures were deposited on slides coated with a 1% agarose buffered solution and imaged as previously described (Diaz et al., 2015). See conditions in Supplemental experimental procedures.

### ChIP-sequencing assay, analysis and fit procedure

ChIP-seq were performed as previously described (Diaz et al., 2017) with minor modifications (Supplemental experimental procedures). Graphing the DNA portion of interest from ChIP-seq data was done using Excel or R softwares. Background levels were determined by normalizing the number of sequence reads between cognate input and IP samples. Data plots superimposed with power law equation were normalized after background subtraction and set to the value of 1 at the last bp of the 10^th^ repeat of *parS*_F_.

### Western immunoblotting

The determination of ParB_F_ relative intracellular concentrations and antibody purifications were performed as described (Diaz et al., 2015). When indicated, samples were diluted in DLT1215 extract to keep constant the total amount of proteins.

### EMSA and proteins purification

EMSA were performed as described (Bouet et al., 2007) in the presence of sonicated salmon sperm DNA as competitor (100 mg.ml^−1^), using 1 nM radiolabeled 144-bp DNA probe containing a single *parS*_F_ site generated by PCR. ParB_F_ and ParB_F_-3R* proteins were purified as previously described (Ah-Seng et al., 2009).

### FRAP and FLIM assays

Cells, grown in mid-exponential phase, were subjected to laser-bleaching over 5-9 pixels (Supplemental experimental procedures). Normalization was performed by averaging fluorescence intensity from the three pre-bleached images.

### Accession number

Raw ChIP-sequencing data for *V. cholera* and *E. coli* are available through the GEO repository with the accession numbers GSE114980 and GSE115274, respectively.

## Acknowledgements

We thank the platform GeT-Biopuces (Genopole, Toulouse) for sequencing experiments and bioinformatics analyses, S. Cantaloube (LITC-CBI platform) for microscopy advices. We are grateful to F. Cornet, P. Polard, P. Rousseau, M. Nollmann, I. Junier and members of the team for fruitful discussions and critical reading of the manuscript. We thank C. Lesterlin for sharing the plasmid F1-10, D. Chattoraj for the anti-ParB*_Vc_*_1_ serum and Y. Yamaichi for pSM836 and *V. cholerae* strains.

## Author contributions

Conceptualization, J.Y.B and A.P.; Methodology, J.Y.B., J.C.W., V.A.L. and A.P.; Investigation, R.E.D., J.C.W, A.S., J.R. and J.Y.B.; Formal Analysis, J.C.W., J.D., F.G., J.P. and A.P.; Writing – Original Draft, R.E.D, J.C.W. and J.Y.B.; Writing – Review & Editing, J.Y.B., J.C.W., R.E.D and A.P.; Funding Acquisition, J.Y.B, A.P., J.P., and V.A.L.; Resources, D.L. and F.B.; Supervision, J.Y.B., V.A.L. and A.P.

